# Impact of decision and action outcomes on subsequent decision and action behaviors

**DOI:** 10.1101/2022.01.24.477509

**Authors:** Clara Saleri Lunazzi, David Thura, Amélie J. Reynaud

**Affiliations:** Lyon Neuroscience Research Center – Impact team Inserm U1028 – CNRS UMR5292 – Lyon 1 University

**Keywords:** decision-making, motor control, reward rate, post-outcome adjustment, arm movement

## Abstract

Speed-accuracy tradeoff adjustments in decision-making have been mainly studied separately from those in motor control. In the wild however, animals coordinate their decision and action, freely investing time in choosing versus moving given specific contexts. Recent behavioral studies support this view, indicating that humans trade decision time for movement time to maximize their global rate of reward during experimental sessions. Besides, it is established that choice outcomes impact subsequent decisions. Crucially though, whether and how a decision also influences the subsequent motor behavior, and whether and how a motor error influences the next decision is unknown. Here we address these questions by analyzing trial-to-trial changes of choice and motor behaviors in healthy human participants instructed to perform successive perceptual decisions expressed with reaching movements whose duration was either bounded or unconstrained in separate tasks. Results indicate that after a bad decision, subjects who were not constrained in their action duration decided more slowly and more accurately. Interestingly, they also shortened their subsequent movement duration by moving faster. Conversely, we found that movement errors not only influenced the speed and the accuracy of the following movement, but those of the decision as well. If the movement had to be slowed down, the decision that precedes that movement was accelerated, and vice versa. Together, these results indicate that from one trial to the next, humans are primarily concerned about determining a behavioral duration as a whole instead of optimizing each of the decision and action speed-accuracy trade-offs independently of each other.

## Introduction

Choosing one action among several options and executing that action are usually considered as two distinct functions, most often studied separately from each other (e.g. Franklin & Wolpert, 2011; Ratcliff et al., 2016). However, recent behavioral studies indicate that decision and action show a high level of integration during goal-oriented behavior (Choi et al., 2014; Cos et al., 2011; Haith et al., 2012; Morel et al., 2017; Shadmehr et al., 2010, 2019; Shadmehr & Ahmed, 2020; Yoon et al., 2018). For example, human subjects decide faster and less accurately to focus on their actions when the motor context in which a choice is made is demanding (Reynaud et al., 2020). Similarly, when the temporal cost of a movement is significantly larger than usual, humans often reduce the duration of their decisions to limit the impact of these time-consuming movements (Saleri Lunazzi et al., 2021). Conversely, if the sensory evidence guiding the choice is weak and the deliberation takes time, humans and monkeys shorten the duration of the movement expressing that choice (Thura, 2020; Thura et al., 2014). Individuals thus seem to be primarily concerned about determining a global behavior duration rather than optimizing decision and action durations separately, even if the resulting decision or movement accuracy must slightly suffer. This “holistic-heuristic” policy may serve what matters the most for decision-makers during successive decisions between actions, the rate of reward (Balci et al., 2011; Carland et al., 2019; Thura, 2021).

Importantly, most of the adjustments mentioned above occur between blocks of tens to hundreds of trials, depending on stable contexts favoring a fixed movement or decision speed-accuracy trade-off. But can these adjustments also occur on shorter time scales, from trial to trial, depending on local decisional and motor performance?

Indeed, performance history is known to exert a large influence on subsequent behavior (e.g. Danielmeier & Ullsperger, 2011; Jentzsch & Dudschig, 2009; Urai et al., 2019). The most well-known post-outcome adjustment is a reduction of behavior speed after committing an error, namely post-error slowing (PES). PES is sometimes accompanied by changes in accuracy, although conditions leading to PES-related increase or decrease of accuracy are still unclear (Danielmeier & Ullsperger, 2011; Fievez et al., 2022). Notably, post-outcome adjustments have been mostly described as the effect of a choice on the decisional performance in the following trial (Dutilh et al., 2012; Laming, 1979; Rabbitt & Rodgers, 1977; Thura et al., 2017; Urai et al., 2019), but the influence of a movement outcome on the motor performance in the following trial did not receive the same attention (Ceccarini & Castiello, 2018). Moreover, the consequences of either a decision or a motor outcome on *both* subsequent decision and movement have never been investigated. These are important questions to address in order to further evaluate the level of integration of the decision and the action functions during goal-directed behavior.

In the present report, we aim at investigating the consequences of *a decision outcome* on the next trial decision *and* motor performance. We also aim at analyzing the effect of *a motor outcome* on the next trial decision *and* motor performance. Because we make the hypothesis that humans decide and act in a “holistic-heuristic” way, we predict that any adjustment due to a decision or a motor outcome will be shared and integrated across the decision and the movement in the next trial. This hypothesis also predicts that the integrated post-outcome adjustments will depend on the capacity of the subject to “freely” share decision time for action time, and vice versa, if needed.

To test this hypothesis, we analyzed datasets from two recent studies of our group during which human subjects made successive perceptual decisions between actions. In the first experiment (Reynaud et al., 2020; Thura, 2020), participants could invest up to 3s in the decision process and had up to 800ms to execute the reaching movement expressing a choice. In the other experiment (Saleri Lunazzi et al., 2021), the decision component of the task was similar but reaching duration was strictly bounded. By analyzing changes of several decision and motor parameters from one trial to the next, we found multiple context-dependent post-decision and post-movement outcome adjustments of both subsequent decision and motor speed-accuracy tradeoffs.

## Material and methods

### Participants

Two groups of healthy, human subjects participated in the two experiments described in this report. Twenty subjects (ages: 20-41; 16 females, 4 males; 18 right-handed) performed the free-movement duration (FMD) task and thirty-one other subjects (ages: 18-36; 20 females, 11 males; 29 right-handed) performed the constrained-movement duration (CMD) task. All gave their consent orally before starting the experiment. The ethics committee of Inserm (IRB00003888) approved the protocol on March 19^th^ 2019. Each participant performed two experimental sessions of the same task. They received monetary compensation for completing each session (either 40 € for the FMD task or 30 € for the CMD task).

### Datasets

The decision and motor behaviors of these subjects have been described in three recent publications reporting the effects of the decisional context on movement properties (Thura, 2020) and the effects of the motor context on decision strategies (Reynaud et al., 2020; Saleri Lunazzi et al., 2021). In these reports, subjects' behavioral adjustments are described either within a given trial (i.e. the relation between a decision duration and the duration of the movement produced to express that decision) or between specific conditions designed to set stable decision or motor speed-accuracy contexts in blocks of tens of trials. Here, we aim at describing adjustments of subjects' behavior from trial to trial, depending on their decision and/or motor performance.Correct and “too slow” movements executed in the CMD task are compared in the bottom right panel. Correct and “too fast” movements executed in the CMD task are compared in the bottom left panel.

### Setup and tasks

The experimental apparatus (figure 1A), identical in the two tasks, as well as visual displays (figure 1B), are detailed and illustrated in the previous publications mentioned above (Reynaud et al., 2020; Saleri Lunazzi et al., 2021; Thura, 2020). The subjects sat in an armchair and made planar reaching movements using a handle held in their dominant hand. A digitizing tablet (GTCO CalComp) continuously recorded the handle horizontal and vertical positions (100 Hz with 0.013 cm accuracy). Target stimuli and cursor feedback were projected by a LCD monitor onto a half-silvered mirror suspended 26 cm above and parallel to the digitizer plane, creating the illusion that targets floated on the plane of the tablet. Participants were faced with a visual display consisting of three blue circles (the decision circles) placed horizontally at a distance of 6 cm from each other. In the central blue circle, 15 tokens were randomly arranged. Positioned below, three black circles, organized horizontally as well, defined the movement targets. The central black circle radius was 0.75 cm. The size and location of the lateral black circles could vary in blocks of trials depending on the task. In the free-movement duration (FMD) task, that size was set to be either 0.75 or 1.5 cm of radius, and distance from the central circle was varied to be either 6 or 12 cm (as mentioned above, effects of target size/position on subjects’ behavior are not included in the present report). In the constrained-movement duration (CMD) task, we analyzed trials for which the target size was set to be 1.5 cm of radius and distance from the central circle was set to 6 cm.

**Figure 1.**
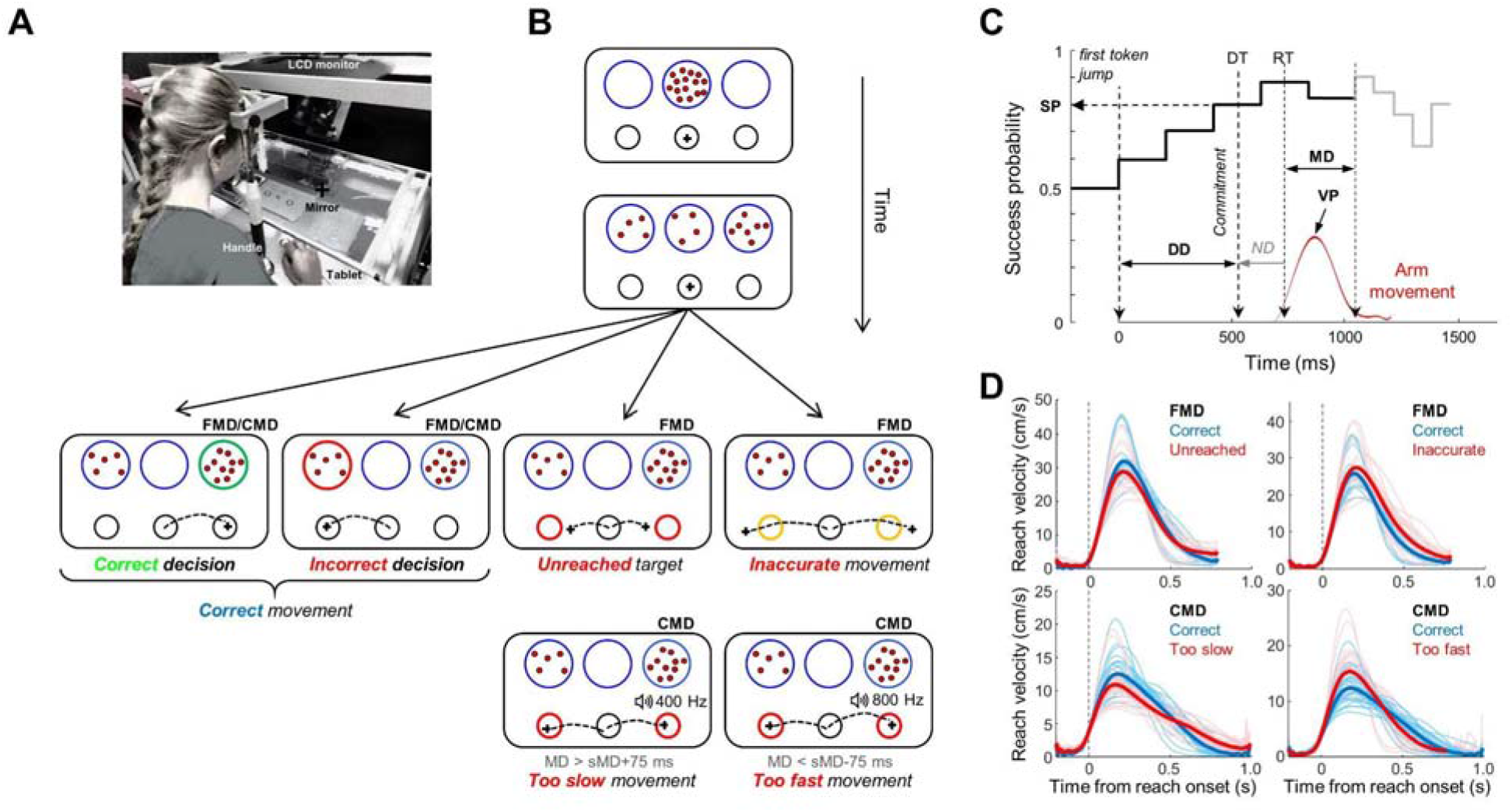
Methods. A. Experimental apparatus, identical in both the FMD and CMD tasks. B. Time course of a trial in the decision task. Tokens jump one-by-one from the central decision circle to one of the two lateral ones. Subjects move a cursor from a central movement target to one of the two lateral ones to express their choice. All the decision and action outcomes are illustrated in the bottom panels (please refer to the main text for details). MD: Movement duration; sMD: Spontaneous movement duration. C. Temporal profile of success probability (SP) in one example trial of the decision task. At the beginning of the trial, each target has the same success probability (0.5). When the first token jumps into one of the two potential targets (the most leftward vertical dotted line), the success probability of that target increases to ~0.6. Success probability then evolves with every jump. Subjects execute a reaching movement (red trace) to report their choice. Kinematic data allow to compute movement duration (MD) and movement peak velocity (VP). Non-decisional (ND) delays, determined in a separate reaction time task, allow to estimate decision duration (DD) and success probability (SP) at decision time. Only 10 out of 15 jumps are illustrated on this SP profile. D. Average reach velocity profiles aligned on reaching movement onset. Correct and “unreached” movements executed in the FMD task are compared in the top left panel; Correct and “inaccurate” movements executed in the FMD task are compared in the top right panel.

In both the FMD and the CMD tasks (figure 1B), implemented by means of LabView 2018 (National Instruments), subjects initiated a trial by holding the handle into the black central circle (starting position) for 500ms. Tokens then started to jump, one by one, every 200ms, in one of the two possible lateral blue circles. Subjects had to decide which of the two lateral blue circles would receive the majority of the tokens at the end of the trial. They reported their decisions by moving the lever into the lateral movement target corresponding to the side of the chosen decision circle. Crucially, participants were allowed to make and report their choice at any time between the first and the last token jump. Once a target was reached, the remaining tokens jumped more quickly to their final circles (figure 1C, gray line), implicitly encouraging subjects to decide before all tokens had jumped to save time and increase their rate of reward at the session level. In the FMD task, tokens could speed up either a lot (a jump every 50ms) or a little (a jump every 150ms) in given blocks of trials. These block-related effects are not included in the present report. In the CMD task, the remaining tokens jumped every 50ms.

In the free-movement duration (FMD) task, subjects had up to 800ms to reach a target and report their choices. If no target was reached within 800ms, trials were classified as “unreached” trials, regardless of the direction of the movement with respect to the starting position. If the subject reached a target but failed to stop in it within 800ms, the trial was classified as “inaccurate” trial, regardless of the choice made, correct or incorrect (figure 1B).

In the constrained-movement duration (CMD) task, participants were instructed to reach a target within a 75-ms time interval around their spontaneous mean movement duration, computed in separate and dedicated trials (please see Saleri Lunazzi et al., 2021 for details). Consequently, if for a given subject we estimated a mean spontaneous reaching duration of 400ms for the 6 cm long movements, then this subject had to report each of her/his choices by executing a movement whose duration was strictly bounded between 325 and 475ms. In this CMD task, a trial was thus considered as a movement error trial when the subject did not meet these temporal constraints, even if the correct decision was made. We distinguished either “too slow” and “too fast” movement errors (figure 1B).

At the end of each trial of both tasks, a visual feedback about decision success or failure (the chosen decision circle turning either green or red, respectively) was provided to the subject after the last token jump, assuming a correct movement. In the FMD task, a movement error was indicated by visual feedback. The chosen movement target turned orange in “inaccurate” trials, the two movement targets turned red in “unreached” trials. In the CMD task, a movement error was indicated by both a visual and a 500ms audio feedback (both movement targets turned red and an 800 or 400 Hz sound indicating that the movement was too fast or too slow, respectively, was played). Subjects had to make a specific number of correct trials (either 320 trials in the FMD task or 160 trials in the CMD task), indirectly motivating them to optimize successes per unit of time.

Finally, subjects also performed in each of the two sessions of both tasks a simple delayed-reaching task (DR task, 100 trials for subjects who performed the FMD task and 20 trials for subjects who performed the CMD task). This DR task was identical to the choice task described above, except that there was only one lateral decision circle displayed at the beginning of the trial (either at the right or at the left side of the central circle with 50% probability). All tokens moved from the central circle to this unique circle at a GO signal occurring after a variable delay (1000 ± 150ms). The DR task was used to estimate the sum of the delays attributable to response initiation (i.e. non-decision delays).

### Subsets of trials based on decision and movement outcomes

We first defined three subsets of trials common to both tasks (FMD and CMD), based on decision or movement outcomes: (1) “Correct decision” trials, when the subject chose the correct target and reported her/his choice with a correct movement; (2) “Incorrect decision” trials, if the participant chose the incorrect target with a correct movement. Note that for these two subsets, bad movement trials are excluded because no feedback was provided to the subject to indicate whether or not she/he chose the correct target. Instead, a salient feedback was provided at the end of the trial to indicate the movement error (see above and figure 1B); (3) “Correct movement” trials, when the subject adequately reached the correct or the incorrect target.

We defined two other subsets of trials based on movement errors in the FMD task specifically: (1) “Unreached” trials, when the subjects failed to reach a target (correct or incorrect) before the end of the movement duration deadline (800ms); (2) “Inaccurate” trials, when the subjects reached a target (correct or incorrect) but failed to stop in it.

Finally, two subsets of trials were defined based on movement errors in the CMD task specifically: (1) “Too fast movement” trials and (2) “too slow movement” trials, when the subjects reached a target (correct or incorrect) before the minimum instructed duration time and after the maximum instructed duration time, respectively.

### Data analysis

Data were analyzed off-line using custom-written MATLAB (MathWorks) and R (https://www.r-project.org/) scripts. Reaching horizontal and vertical positions were first filtered using polynomial filters and then differentiated to obtain a velocity profile. Onset and offset of movements were then determined using a 3.75 cm/s velocity threshold. Reaching movement duration (MD), peak velocity (VP) and amplitude (Amp) were respectively defined as the duration, the maximum velocity value and the Euclidean distance between these two events (figure 1C). Reaching movement accuracy was defined as the Euclidian distance separating the target center from the movement endpoint location (CED).

Decision duration (DD) was computed as the duration between the first token jump and the time at which subjects committed to their choice (figure 1C). To estimate this commitment time in each trial, we detected the time of movement onset as mentioned above, defining the subject's reaction time, and subtracted from it her/his mean sensory-motor delays estimated based on her/his reaction times in the DR task performed the same day and in the same condition.

To assess the influence of sensory evidence on subjects' choices, we computed the success probability profile of each trial experienced by participants with respect to the chosen target, as well as their decision success probability (SP) at the time of commitment time (figure 1C), using Equation 1. For instance, for a total of 15 tokens, if at a particular moment in time the target chosen by the subject contains N_chosen_ tokens, whereas the other target contains N_other_ tokens, and there are NC tokens remaining in the center, then the probability that the chosen target will ultimately be the correct one, i.e. the subject's success probability (SP) at a particular time is as follows:

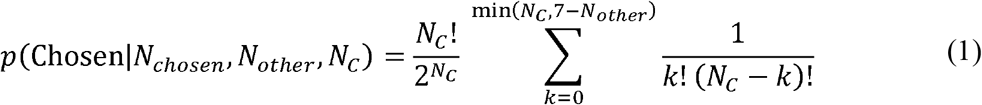

To ensure that the difficulty of decisions was homogeneous among subjects and experimental conditions, we controlled the sequence of trials experienced by each participant in each session of both tasks. Especially, we interspersed among fully random trials (~20% of the trials in which each token is 50% likely to jump into the right or the left lateral circle) three special types of trials, easy, ambiguous and misleading, characterized by particular temporal profiles of success probability. Subjects were not told about the existence of these trials. Please refer to Reynaud et al., 2020 and Saleri Lunazzi et al., 2021 for a detailed description of these trial types and their proportions in the FMD and CMD tasks.

To assess the impact of the outcome of each trial *i* on the decision and motor behavior of trial *i+1,* we calculated the difference of movement velocity peak (ΔVP), duration (ΔMD), amplitude (ΔAmp), accuracy (ΔCED), and the difference of decision duration (ΔDD) and success probability (ΔSP) between them (e.g. ΔVP = *VP_i+1_* – *VP_i_*. We then calculated for each subject the average of each variable with respect to trial *i* outcome.

### Statistics

To determine whether the behavioral adjustment from one trial to the following (ΔVP, ΔMD, ΔAmp, ΔCED, ΔDD and ΔSP) differs significantly from 0 in the different outcome conditions at the population level, we used one-sample Wilcoxon signed rank tests. A Levene's test was used to test if the distributions of the post-correct and post-error decision and motor variables have equal variances. Pearson's correlation tests were used to directly investigate the relationship between motor (ΔVP, ΔMD, ΔAmp, ΔCED) and decision (ΔDD, ΔSP) adjustments following different outcomes. For all statistical tests, the significance level is set to 0.05. Unless stated otherwise, data are reported as medians across the population. To estimate the difference between the average success probability profiles of two trial subsets (e.g. correct decision trials versus post-correct decision trials), we computed the distance between the two profiles (1 and 2) from token jump (*j*) #1 to #15 as the following chi-squared metric:

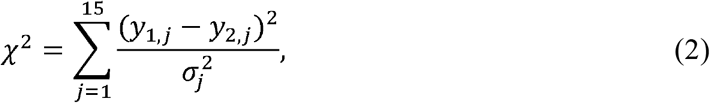

where *y_1_* and *y_2_* are the two SP profiles averaged across subjects, and 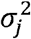 is the mean squared variance of the SP profiles, such as 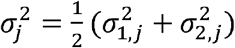.

## Results

### Effect of a decision outcome on the next decision and on the next movement in the FMD task

We first describe the impact of the decision outcome (correct or incorrect choice) on participants' subsequent decisional behavior when the motor temporal constraints were low (FMD task). As shown in figure 2A, subjects' decision duration was significantly increased compared to a previous incorrect decision (median ΔDD=+70.6ms, Wilcoxon signed rank test, Z=2.4, p=0.015). This slowdown of decision-making was observed despite that trials following an incorrect choice were easier, as can be seen on the averaged success probability (SP) profiles of the two trial subsets (**χ**^2^=2119, inset in figure 2A, top right panel; Suppl. figure 1 illustrates the SP profiles of the same trials computed with respect to the correct target). As a consequence, subjects' SPs at decision time were increased following incorrect decisions (ΔSP=+0.09, Z=3.9, p<0.001). By contrast, no significant difference of decision duration (ΔDD=-2.8ms) was observed following a correct decision. Together, this first analysis demonstrates that most subjects used a post-error slowing strategy to decide in this task, as can be seen when decision durations following either a correct or a bad choice are directly compared (suppl. figure 2). Interestingly, subjects did not adjust their decision duration following a correct trial despite that these trials were on average slightly more difficult (**χ**^2^=207, inset in figure 2A, bottom right panel). Participants’ success probability thus slightly decreased after a correct choice (ΔSP=-0.02, Z=-3.9, p<0.001), indicating that they committed to a decision with less sensory evidence after a correct trial.

**Figure 2.**
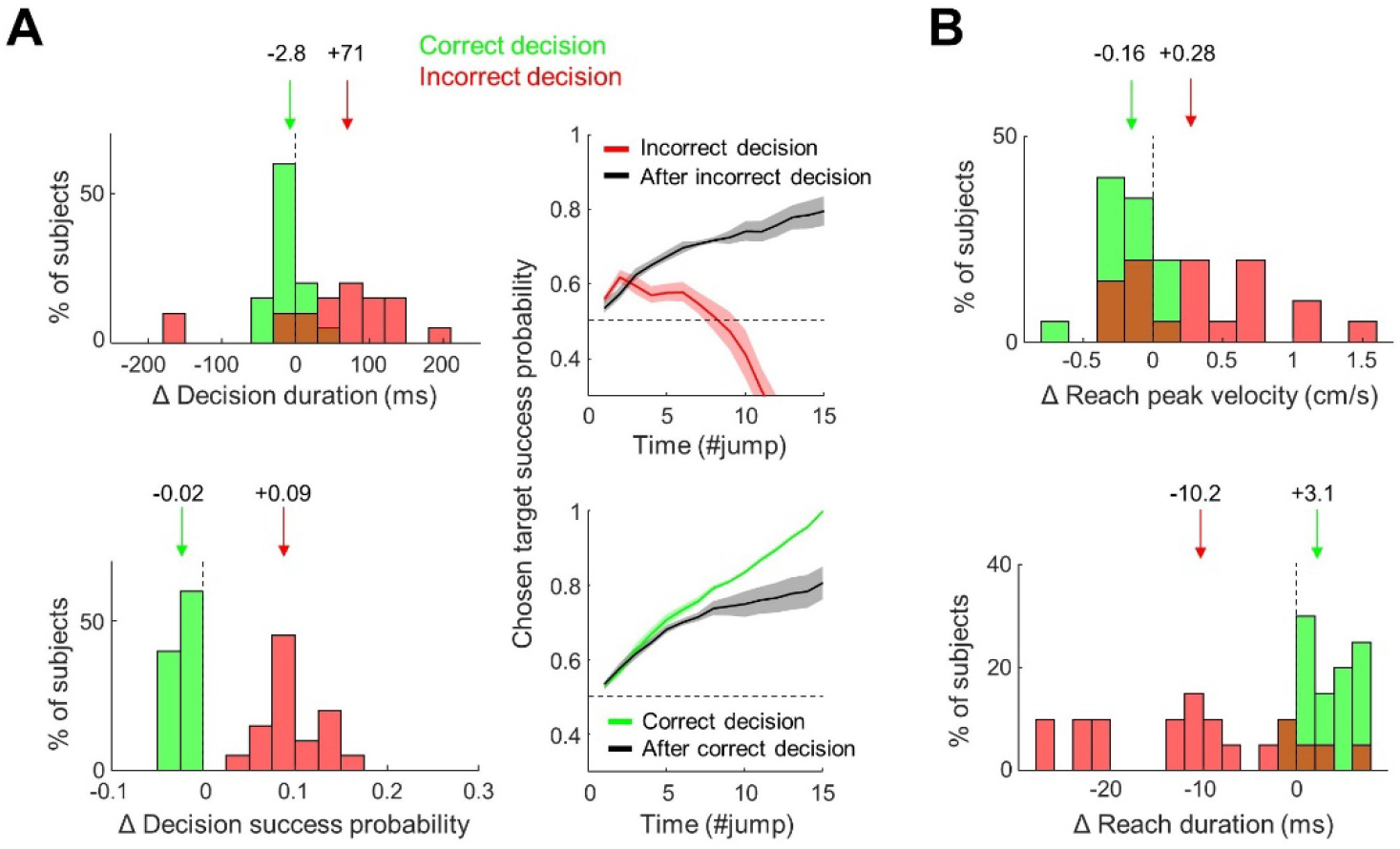
Effect of a decision outcome on the next decision and on the next movement in the FMD task. A. Left panels: Distribution and comparison of decision duration (top) and success probability (bottom) adjustments depending on the decision outcome in the previous trial (after a correct trial in green or after an incorrect decision in red). Arrows mark the population medians whose values are reported above. The dotted black line indicates zero difference between the trial *i* and *i+1.* If Δ is positive, there is a post-outcome increase for a given metric X (*X_i+1_* – X*_i_* >0) whereas a negative Δ value indicates a decrease of this metric. Right panel, top: Comparison of the average ± SD success probability profiles between trials whose decision was incorrect (red solid line) and trials following an incorrect choice (black dotted line), computed across subjects with respect to the target they chose. Right panel, bottom: same comparison between correct decision trials (green solid line) and trials following a correct decision (black solid line). B: Same analysis as in A, left panels, for the post-decision outcome adjustments computed for movement peak velocity (top) and duration (bottom).

We next investigate whether or not a decision outcome also impacts motor behavior. We found that following incorrect decisions, subjects made overall faster movements (ΔVP=+0.28 cm/s, Z=2.4, p=0.01), thus reducing their reaching duration (ΔMD=-10.2ms, Z=3.5, p<0.001, figure 2B) despite a decrease of amplitude (ΔAmp=-0.08 cm, Z=-3, p=0.002, suppl. figure 3). We also observed at the population level an increase of movement inaccuracy after an incorrect choice (ΔCED=+0.06 cm, Z=3.1, p=0.002), but this effect was also observed for trials following correct decisions (ΔCED=+0.06 cm, Z=3.9, p<0.001, suppl. figure 3).

### Effect of a movement outcome on the next decision and on the next movement in the FMD task

In the free-movement duration (FMD) task, we distinguished two types of movement error: “Inaccurate” trials, when a target was reached but subjects failed to stop in it, and “unreached” trials, when the subject failed to reach a target before the movement duration deadline (800ms in the FMD task). “Inaccurate” movements were thus on average faster (26.8 vs 26 cm/s), larger in amplitude (9.2 vs 8.7 cm) and longer (557 vs 530ms) compared to correct movements (figure 1D, top right panel). As expected, subjects corrected these inaccurate movements in the following trial (ΔCED=-0.19 cm, Z=-2.6, p=0.01) by decreasing their reaching velocity peak (ΔVP=-1.7 cm/s, p=0.001, Z=-3.2, figure 3A, top panels). Participants also reduced their movement amplitude (ΔAmp=-1.1 cm, Z=-3.9, p<0.001) and duration (ΔMD=-40ms, Z=-3; p=0.002) in trials following an inaccurate movement (suppl. figure 4A).

**Figure 3.**
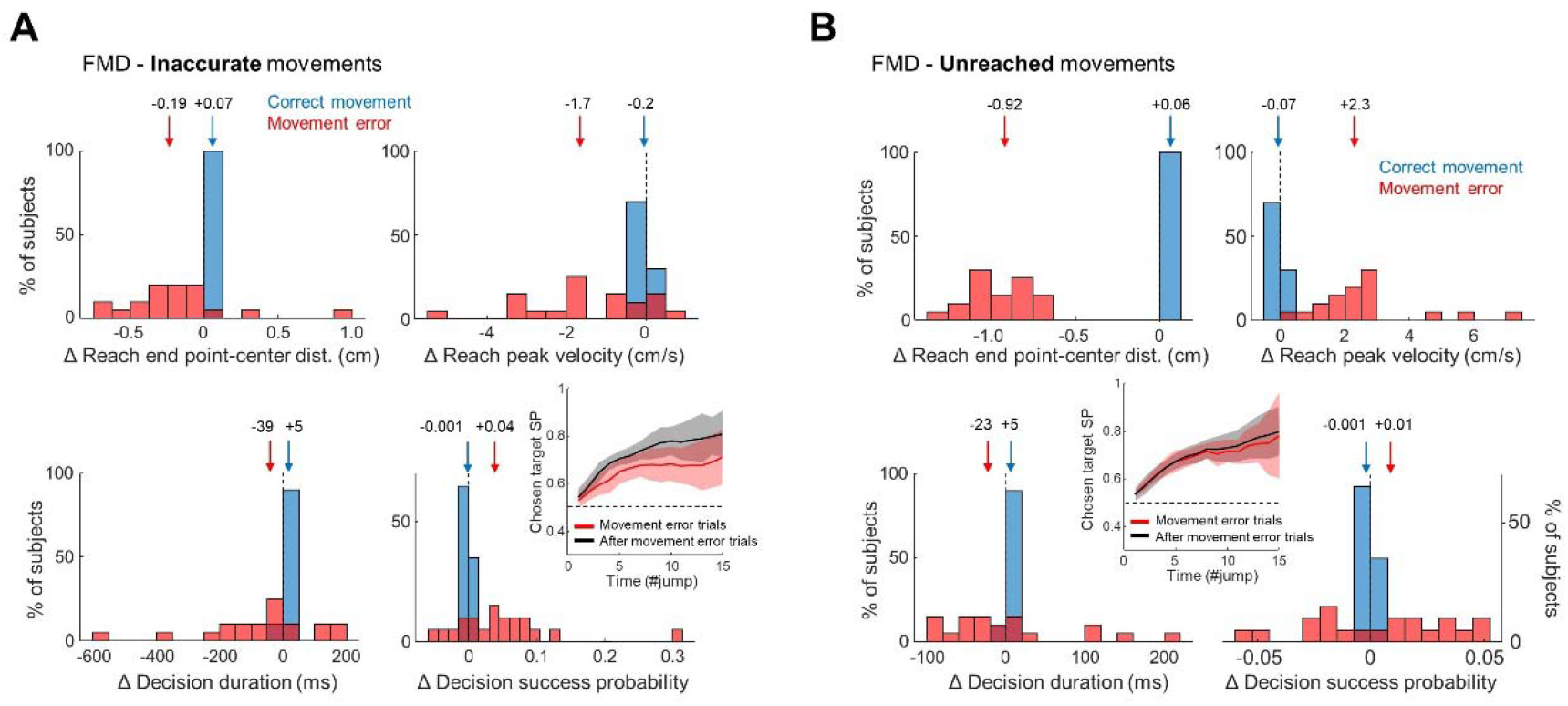
Effect of movement errors on subsequent motor and decision behaviors in the FMD task. A. Top: Distribution and comparison of reaching movement end-point center distance (left) and peak velocity (right) adjustments depending on the movement outcome in the previous trial (after a correct movement in blue and after an “inaccurate” movement in red). Bottom: Distribution and comparison of decision duration (left) and success probability (right) adjustments depending on the movement outcome in the previous trial (after a correct movement in blue and after an “inaccurate” movement in red). The inset illustrates the average ± SD success probability profiles of inaccurate movement trials (red) and post-inaccurate movement trials (black), computed across subjects. B. Same as A for the “unreached” trials.

As illustrated in the bottom panels of figure 3A, population mean decision durations and success probabilities are much variable and distributed in trials following an inaccurate movement compared to trials following a correct movement (SD=182 versus 4.6ms, respectively; Levene's test, F=21.7, p<0.0001). In terms of medians, decision durations following an inaccurate movement were overall shorter compared to trials for which a movement was inaccurate, but this difference is not significant (ΔDD=-39ms). We also observed a slight but significant increase of decision success probability (ΔSP=+0.04, Z=2.8, p=0.005) following inaccurate movements, possibly because of the slightly higher SP profile of trials following inaccurate movements compared to the error movement trials (**χ**^2^=21.8, figure 3A, bottom right panel). To directly assess the relationship between motor and decision adjustments due to inaccurate movements, we computed linear regressions between all differences of motor (ΔVP, ΔMD, ΔCED, ΔAmp) and decision (ΔDD, ΔSP) metrics. We found a significant negative correlation between ΔMD and ΔDD (Pearson correlation, R=-0.62, p=0.003) and a significant positive correlation between ΔMD and ΔSP (R=0.66, p=0.003), indicating that subjects who decreased their subsequent reaching duration the most after an inaccurate movement increased their subsequent decision duration and decrease their subsequent decision success probability the most as well (suppl. figure 4B).

While various reasons could lead to “unreached” trials, we noticed that overall, movements in these trials were on average slower compared to correct movements (figure 1D, top left panel). Following these unreached trials, subjects significantly increased their movement accuracy in the next trial (ΔCED=-0.92 cm, Z=-3.9, p<0.001) by increasing their reaching velocity peak (ΔVP=+2.3 cm/s, Z=3.9, p<0.001, figure 3B, top panels). They also increased their reaching amplitude (ΔAmp=+1 cm, Z=3.9, p<0.001) and duration (ΔMD=+20ms, Z=2.9, p=0.004) compared to the previous erroneous trials (suppl. figure 4C).

After an “unreached” movement, we did not observe significant adjustments of the decisional behavior in the next trial at the population level (ΔDD=-23ms, ΔSP=+0.01), although distributions of mean decision durations and success probabilities are broader in trials following an unreached movement compared to trials following a correct movement (SD= 83 versus 4.6ms; Levene's test, F=25, p<0.0001 figure 3B, bottom panels). We found however a significant positive correlation between the adjustment of peak velocities (ΔVP) following an unreached movement trial and the adjustment of decision durations (ΔDD) in the same condition (R=0.48, p=0.04, suppl. figure 4D), indicating that participants who increased their movement speed the most after an unreached movement trial also increased their decision duration the most.

### Post-outcome adjustments of decision and motor behaviors in the CMD task

The previous paragraphs describe behavioral adjustments of subjects performing the free-movement duration (FMD) task. In the following lines, we report the same analyses applied on a dataset collected in the constrained movement duration (CMD) task. In this task, the duration of a movement executed to report a choice was strictly bounded (see methods), increasing the difficulty of the motor aspect of the task. In the FMD task, the average percentage of error trials was 25%, among which 18% of decisional errors and 7% of movement errors. In the CMD task however, about 50% of trials were unsuccessful, with only 12% of decisional errors but 38% of movement errors.

We first assessed whether a decision outcome influenced the subsequent decision behavior in the CMD task. As shown in figure 4A, the slowdown of decisions observed following incorrect choices in the FMD task was not found in the CMD task (ΔDD=-5ms). Subsequent decision success probabilities were increased following incorrect decisions (ΔSP=+0.12, Z=4.8, p<0.001), an adjustment likely due to post-incorrect decision trials that were easier compared to incorrect decision trials **χ**^2^=767, inset in figure 4A, right panel). Despite that a decision outcome did not influence the next decision in the CMD task, we observed that following incorrect choices, participants increased their reaching velocity peak (ΔVP=+0.24 cm/s, Z=3, p=0.003), leading to a decrease of movement duration (ΔMD=-13ms, Z=-3.4, p<0.001, figure 4B).

**Figure 4.**
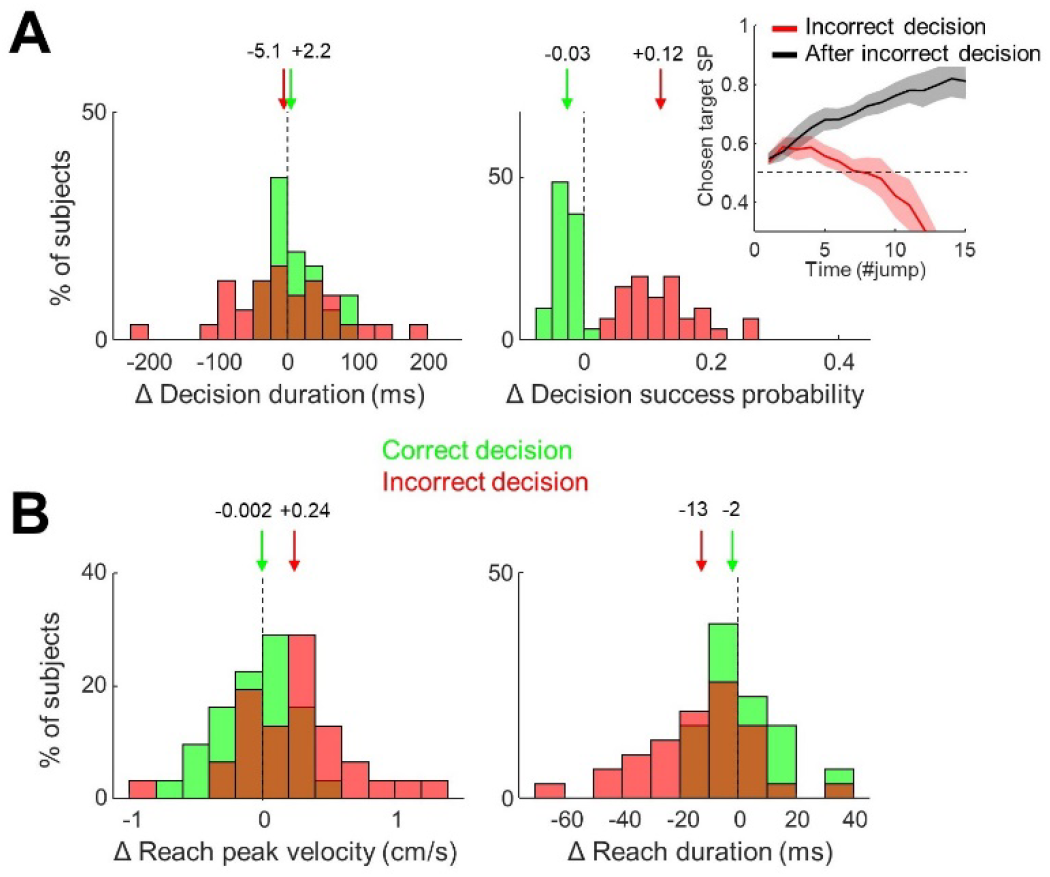
Effect of decision outcomes on subsequent decision and motor behaviors in the CMD task. A. Distribution and comparison of decision duration (left) and success probability (right) adjustments depending on the decision outcome in the previous trial (after a correct decision in green and after an error decision in red). The inset illustrates the average ± SD success probability profiles of incorrect decision trials (red) and post-incorrect decision trials (black), computed across subjects. B: Distribution and comparison of reaching peak velocity (left) and duration (right) adjustments depending on the decision outcome in the previous trial (same conventions as in A).

In the last two paragraphs, we investigate the consequences of a movement error in the CMD task, i.e. too fast or too slow movements, on subjects' behavior in the next trial. As expected, following too fast movements, participants significantly decreased their reaching velocity peak (ΔVP=-3.1 cm/s, Z=-4.8, p<0.001), leading to an increase of movement duration (ΔMD=+62ms, Z=4.8, p<0.001, figure 5A, top panels). This adjustment was accompanied by a decrease of amplitude (ΔAmp=-0.35 cm, Z=-4.8, p<0.001) and a decrease of accuracy (ΔCED=+0.2 cm, Z=4.8, p<0.001, suppl. figure 5A). The duration of decisions at the population level was significantly decreased following too fast movements in the CMD task (ΔDD=-79ms, Z=-3.7, p<0.001, figure 5A, bottom left panel). Crucially, this adjustment is not due to a difference of decision difficulty between the two trial subsets (**χ**^2^ =1.2, inset in figure 5A, bottom right panel), and no variation of decision success probability (SP) was observed following too fast movement trials.

**Figure 5.**
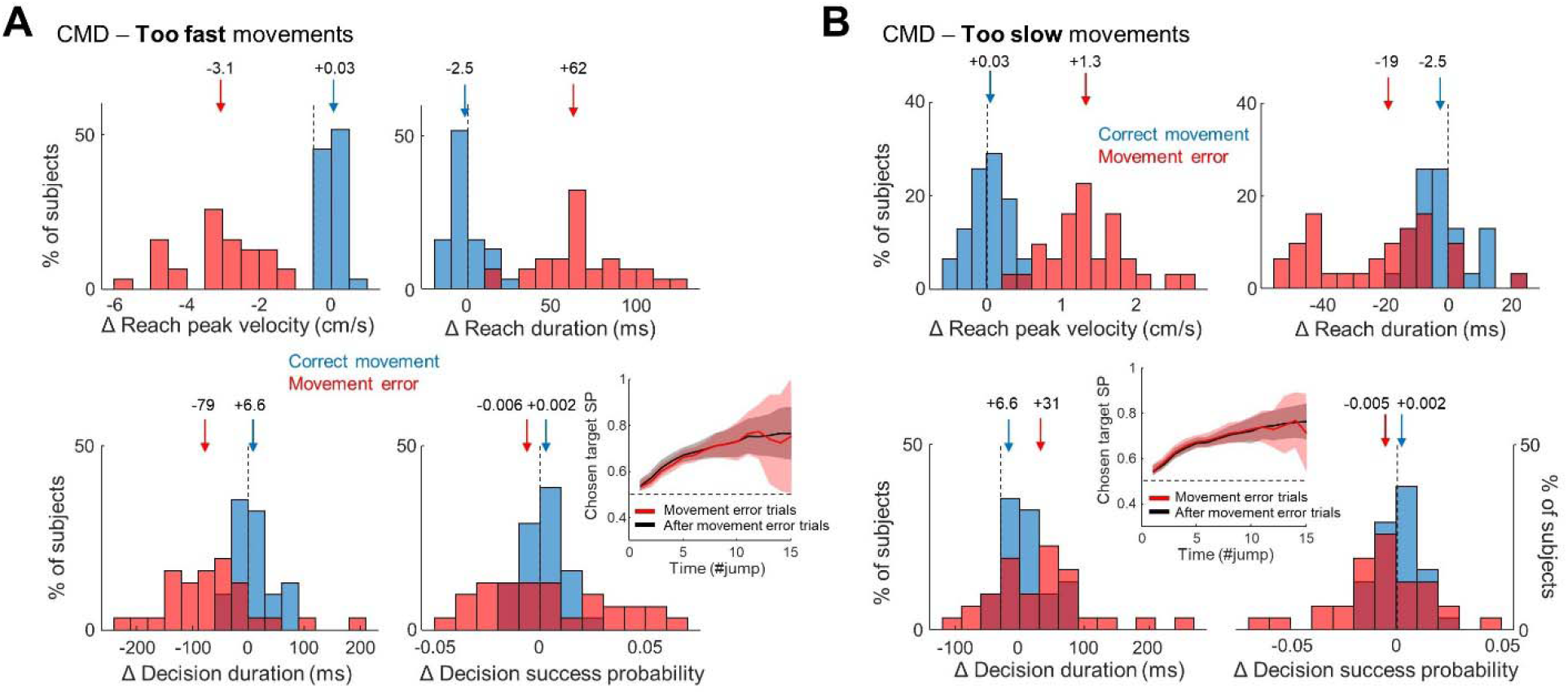
Effect of movement errors on subsequent motor and decision behaviors in the CMD task. A. Top: Distribution and comparison of reaching movement peak velocity (left) and duration (right) adjustments depending on the movement outcome in the previous trial (after a correct movement in blue and after a too fast movement in red). Bottom: Distribution and comparison of decision duration (left) and success probability (right) adjustments depending on the movement outcome in the previous trial (same convention as above). The inset illustrates the average ± SD success probability profiles of too fast movement trials (red) and post-too fast movement trials (black), computed across subjects. B. Same as A for too slow movement trials in the CMD task.

After too slow movements, participants unsurprisingly increased their movement velocity peak (ΔVP=+1.3 cm/s, Z=4.8, p<0.001), which reduced reaching durations (ΔMD=-19ms, Z=-4.4, p<0.001, figure 5B, top panels). Movement amplitude (ΔAmp=+0.3 cm, Z=4.8, p<0.001) and accuracy (ΔCED=-0.29 cm, Z=-4.8, p<0.001) were also significantly increased (suppl. figure 5B). Notably, following too slow movements, the duration of decisions was in this case significantly increased (ΔDD=+31ms, Z=2.1, p=0.038), despite no difference between the SP profiles in the tow trial subsets (**χ**^2^ =0.5), without a significant modulation of decision SP (figure 5B, bottom panels).

## Discussion

In the present study we first observed that in the free-movement duration (FMD) task, most subjects slowed down their choices following a decision error. Post-(decision) error slowing (PES) is a phenomenon commonly reported in the literature (Danielmeier & Ullsperger, 2011; Jentzsch & Dudschig, 2009; Laming, 1979; Purcell & Kiani, 2016; Rabbitt & Rodgers, 1977; Thura et al., 2017), even if post-error speeding has been described as well (e.g. King et al., 2010). PES is often interpreted as an error-induced increase in response caution that allows one to improve subsequent performance Interestingly, after a correct choice, subjects did not adjust their choice durations, but they committed with less sensory evidence. Because post-correct decision trials were slightly more difficult than correct trials, this result suggests that a successful behavior increased participants' confidence, possibly promoting risk taking (Bandura & Locke, 2003).

The present report reveals the properties of the decision-related PES further by showing that participants did not adjust their decision duration following a bad decision in the constrained-movement duration (CMD) task. To explain this observation, it could first be argued that post-decision error trials were overall easier compared to trials in which a decision error occurred. Although possible, the difference of success probability profiles in the two trial subsets was similar in the FMD and CMD tasks, suggesting another reason for the lack of PES in the CMD task. Alternatively, it is known that PES partly depends on error frequency (Notebaert et al., 2009), and participants made more errors in the CMD task compared to the FMD task. However, errors in the CMD task concerned mostly movements, and decision error rates were similar in the two tasks. We thus believe that the lack of decision-related PES in the CMD task primarily relates to the strict duration constraints imposed on movements in this task (see below).

We also observed the expected, yet very robust, post-movement error adjustments in participants' motor behavior. Generally, effects of behavior history on subsequent behavior have been investigated by means of cognitive tasks (Dutilh et al., 2012; Notebaert et al., 2009; Rabbitt & Rodgers, 1977), limiting the analysis to pre-movement processes (but see Ceccarini & Castiello, 2018). The present report describes, to our knowledge, the first analysis addressing the impact of decision *and* action outcomes on *both* the decision and action executed in the following trial. This is important because in most everyday life choices, decisions and movements expressing these choices are temporally linked, constituting a continuum separating an event from a potential reward (Cisek, 2007).

We found that after a slower choice made in response to a decision error, movement duration, if unconstrained, is reduced. This result is consistent with recent reports in both human and non-human primates showing that within blocks of trials defined by specific speed-accuracy tradeoff (SAT) properties, long decisions are expressed with vigorous, short movements (Thura, 2020; Thura et al., 2014). We show here that this policy can be established on a shorter time scale, from trial to trial, based on subject's previous trial outcomes.

Conversely, we found that when participants had to correct a bad movement, they not only adequately adjusted their movements in the following trial, but they also altered the decision made in this following trial, prior to the corrected movement expressing that choice. This observation is at first sight consistent with several studies showing that the cost of a movement executed to report a choice influences that choice in a given trial (Burk et al., 2014; Hagura et al., 2017; Marcos et al., 2015). But it actually differs by demonstrating for the first time the ability of humans to preemptively compensate for a movement correction due to a motor error by altering the deliberation process of the post-error trial, before the execution of the corrected movement.

A possible functional interpretation of the reduction of movement duration accompanying a decision-related PES (in the FMD task) is that subjects aimed at compensating the extra time devoted to deliberation by executing faster movements, even if shortening movement duration usually leads to a slight decrease of accuracy. In ecological scenarios, individuals are indeed often free to adjust the time they invest in deciding versus moving, and movements are parametrized following “economic” rules (e.g. Shadmehr et al., 2019), allowing to optimize what matters the most for individuals during successive choices, the rate of reward (Balci et al., 2011; Bogacz et al., 2010; Carland et al., 2019).

In agreement with a reward rate maximization account, when a movement was corrected by increasing or decreasing its duration, most participants decreased or increased their decision duration, respectively, within the same trial. This is consistent with our previous reports in which compensatory effects are described across blocks of tens of trials defined by specific motor SAT constraints (Reynaud et al., 2020; Saleri Lunazzi et al., 2021). This suggests that one can flexibly share temporal resources between the decision and the action processes depending on both global and local contexts, even if these processes must slightly suffer in terms of accuracy (i.e. a good enough, or heuristic, approach, Gigerenzer & Gaissmaier, 2011). According to this mechanism, the absence of decision-related PES when movement duration was strictly bounded would mean that subjects anticipated that they could not compensate for a potential extension of their decision duration following a bad choice during the movement phase, discouraging them to slow down their decisions after a decision error. Intriguingly, they still produced faster and shorter movements after a bad choice, indicating here an adjustment of movement duration that does not depend on the decision determining this movement. It is possible that in this specific task where errors were frequent (~50%), subjects aimed at limiting the waste of time due to an erroneous trial by moving slightly faster in the next trial despite the strict constraints imposed on movement duration.

Taken together, the present results indicate that following both decision and movement errors, humans are primarily concerned about determining a behavioral duration as a whole instead of optimizing each of the decision and action speed-accuracy trade-offs independently of each other, probably with the goal of maximizing their success rate.

## Open practices statement

The data and materials for the two experiments are available upon request. None of the experiments was preregistered.

## Acknowledgements/Funding

The authors have no financial or non-financial interests to declare This work is supported by a CNRS/Inserm ATIP/Avenir grant to David Thura

## References

Balci, F., Simen, P., Niyogi, R., Saxe, A., Hughes, J. A., Holmes, P., & Cohen, J. D. (2011). Acquisition of decision making criteria: Reward rate ultimately beats accuracy. Attention, Perception, & Psychophysics, 73(2), 640–657. https://doi.org/10.3758/s13414-010-0049-7

Bandura, A., & Locke, E. A. (2003). Negative self-efficacy and goal effects revisited. Journal of Applied Psychology, 88(1), 87–99. https://doi.org/10.1037/0021-9010.88.1.87

Bogacz, R., Hu, P. T., Holmes, P. J., & Cohen, J. D. (2010). Do humans produce the speed–accuracy trade-off that maximizes reward rate? Quarterly Journal of Experimental Psychology, 63(5), 863–891. https://doi.org/10.1080/17470210903091643

Burk, D., Ingram, J. N., Franklin, D. W., Shadlen, M. N., & Wolpert, D. M. (2014). Motor Effort Alters Changes of Mind in Sensorimotor Decision Making. PLoS ONE, 9(3), e92681. https://doi.org/10.1371/journal.pone.0092681

Carland, M. A., Thura, D., & Cisek, P. (2019). The Urge to Decide and Act: Implications for Brain Function and Dysfunction. The Neuroscientist, 107385841984155. https://doi.org/10.1177/1073858419841553

Ceccarini, F., & Castiello, U. (2018). The grasping side of post-error slowing. Cognition, 179, 1–13. https://doi.org/10.1016/j.cognition.2018.05.026

Choi, J. E. S., Vaswani, P. A., & Shadmehr, R. (2014). Vigor of Movements and the Cost of Time in Decision Making. Journal of Neuroscience, 34(4), 1212–1223. https://doi.org/10.1523/JNEUROSCI.2798-13.2014

Cisek, P. (2007). Cortical mechanisms of action selection: The affordance competition hypothesis. Philosophical Transactions of the Royal Society B: Biological Sciences, 362(1485), 1585–1599. https://doi.org/10.1098/rstb.2007.2054

Cos, I., Bélanger, N., & Cisek, P. (2011). The influence of predicted arm biomechanics on decision making. Journal of Neurophysiology, 105(6), 3022–3033. https://doi.org/10.1152/jn.00975.2010

Danielmeier, C., & Ullsperger, M. (2011). Post-Error Adjustments. Frontiers in Psychology, 2, 233. https://doi.org/10.3389/fpsyg.2011.00233

Dutilh, G., Vandekerckhove, J., Forstmann, B. U., Keuleers, E., Brysbaert, M., & Wagenmakers, E.-J. (2012). Testing theories of post-error slowing. Attention, Perception & Psychophysics, 74(2), 454–465. https://doi.org/10.3758/s13414-011-0243-2

Fievez, F., Derosiere, G., Verbruggen, F., & Duque, J. (2022). Post-error Slowing Reflects the Joint Impact of Adaptive and Maladaptive Processes During Decision Making. Frontiers in Human Neuroscience, 16, 864590. https://doi.org/10.3389/fnhum.2022.864590

Franklin, D. W., & Wolpert, D. M. (2011). Computational mechanisms of sensorimotor control. Neuron, 72(3), 425–442. https://doi.org/10.1016/j.neuron.2011.10.006

Gigerenzer, G., & Gaissmaier, W. (2011). Heuristic Decision Making. Annual Review of Psychology, 62(1), 451–482. https://doi.org/10.1146/annurev-psych-120709-145346

Hagura, N., Haggard, P., & Diedrichsen, J. (2017). Perceptual decisions are biased by the cost to act. ELife, 6, e18422. https://doi.org/10.7554/eLife.18422

Haith, A. M., Reppert, T. R., & Shadmehr, R. (2012). Evidence for Hyperbolic Temporal Discounting of Reward in Control of Movements. Journal of Neuroscience, 32(34), 11727–11736. https://doi.org/10.1523/JNEUROSCI.0424-12.2012

Jentzsch, I., & Dudschig, C. (2009). Short Article: Why do we slow down after an error? Mechanisms underlying the effects of posterror slowing. Quarterly Journal of Experimental Psychology, 62(2), 209–218. https://doi.org/10.1080/17470210802240655

King, J. A., Korb, F. M., von Cramon, D. Y., & Ullsperger, M. (2010). Post-error behavioral adjustments are facilitated by activation and suppression of task-relevant and task-irrelevant information processing. The Journal of Neuroscience: The Official Journal of the Society for Neuroscience, 30(38), 12759–12769. https://doi.org/10.1523/JNEUROSCI.3274-10.2010

Laming, D. (1979). Choice reaction performance following an error. Acta Psychologica, 43(3), 199–224. https://doi.org/10.1016/0001-6918(79)90026-X

Marcos, E., Cos, I., Girard, B., & Verschure, P. F. M. J. (2015). Motor Cost Influences Perceptual Decisions. PLOS ONE, 10(12), e0144841. https://doi.org/10.1371/journal.pone.0144841

Morel, P., Ulbrich, P., & Gail, A. (2017). What makes a reach movement effortful? Physical effort discounting supports common minimization principles in decision making and motor control. PLOS Biology, 15(6), e2001323. https://doi.org/10.1371/journal.pbio.2001323

Notebaert, W., Houtman, F., Opstal, F. V., Gevers, W., Fias, W., & Verguts, T. (2009). Post-error slowing: An orienting account. Cognition, 111(2), 275–279. https://doi.org/10.1016/j.cognition.2009.02.002

Purcell, B. A., & Kiani, R. (2016). Neural Mechanisms of Post-error Adjustments of Decision Policy in Parietal Cortex. Neuron, 89(3), 658–671. https://doi.org/10.1016/j.neuron.2015.12.027

Rabbitt, P., & Rodgers, B. (1977). What does a man do after he makes an error? An analysis of response programming. Quarterly Journal of Experimental Psychology, 29(4), 727–743. https://doi.org/10.1080/14640747708400645

Ratcliff, R., Smith, P. L., Brown, S. D., & McKoon, G. (2016). Diffusion Decision Model: Current Issues and History. Trends in Cognitive Sciences, 20(4), 260–281. https://doi.org/10.1016/j.tics.2016.01.007

Reynaud, A. J., Saleri Lunazzi, C., & Thura, D. (2020). Humans sacrifice decision-making for action execution when a demanding control of movement is required. Journal of Neurophysiology, 124(2), 497–509. https://doi.org/10.1152/jn.00220.2020

Saleri Lunazzi, C., Reynaud, A. J., & Thura, D. (2021). Dissociating the Impact of Movement Time and Energy Costs on Decision-Making and Action Initiation in Humans. Frontiers in Human Neuroscience, 15, 715212. https://doi.org/10.3389/fnhum.2021.715212

Shadmehr, R., & Ahmed, A. A. (2020). Vigor: Neuroeconomics of Movement Control. The MIT Press. https://doi.org/10.7551/mitpress/12940.001.0001

Shadmehr, R., Orban de Xivry, J. J., Xu-Wilson, M., & Shih, T.-Y. (2010). Temporal Discounting of Reward and the Cost of Time in Motor Control. Journal of Neuroscience, 30(31), 10507–10516. https://doi.org/10.1523/JNEUROSCI.1343-10.2010

Shadmehr, R., Reppert, T. R., Summerside, E. M., Yoon, T., & Ahmed, A. A. (2019). Movement Vigor as a Reflection of Subjective Economic Utility. Trends in Neurosciences, 42(5), 323–336. https://doi.org/10.1016/j.tins.2019.02.003

Thura, D. (2020). Decision urgency invigorates movement in humans. Behavioural Brain Research, 382, 112477. https://doi.org/10.1016/j.bbr.2020.112477

Thura, D. (2021). Reducing behavioral dimensions to study brain–environment interactions. Behavioral and Brain Sciences, 44. https://doi.org/10.1017/S0140525X21000169

Thura, D., Cos, I., Trung, J., & Cisek, P. (2014). Context-dependent urgency influences speed-accuracy trade-offs in decision-making and movement execution. The Journal of Neuroscience: The Official Journal of the Society for Neuroscience, 34(49), 16442–16454. https://doi.org/10.1523/JNEUROSCI.0162-14.2014

Thura, D., Guberman, G., & Cisek, P. (2017). Trial-to-trial adjustments of speed-accuracy trade-offs in premotor and primary motor cortex. Journal of Neurophysiology, 117(2), 665–683. https://doi.org/10.1152/jn.00726.2016

Urai, A. E., de Gee, J. W., Tsetsos, K., & Donner, T. H. (2019). Choice history biases subsequent evidence accumulation. ELife, 8, e46331. https://doi.org/10.7554/eLife.46331

Yoon, T., Geary, R. B., Ahmed, A. A., & Shadmehr, R. (2018). Control of movement vigor and decision making during foraging. Proceedings of the National Academy of Sciences, 115(44), E10476–E10485. https://doi.org/10.1073/pnas.1812979115

