## Supplemental material for "Impact of decision and action outcomes on subsequent decision and action behaviors"

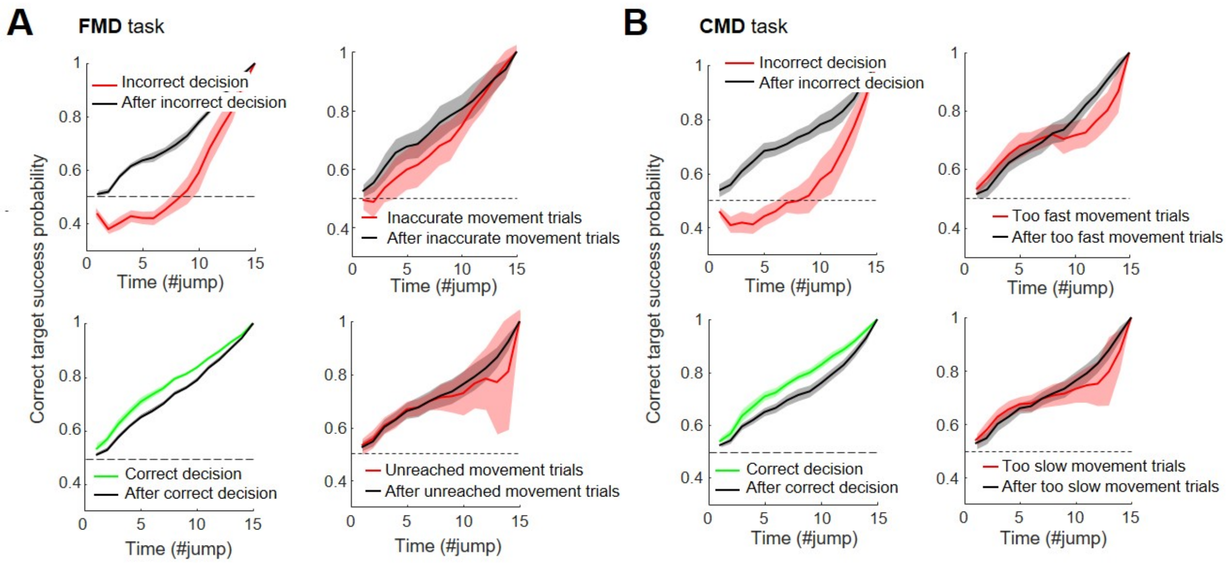


**Supplemental figure 1.** A. Top-left: Comparison of the average ± SD success probability profiles between trials whose decision was incorrect (red solid line) and trials following an incorrect choice (black dotted line), computed across subjects with respect to the correct target, in the FMD task. Bottom-left: Same comparison between correct decision trials (green solid line) and trials following a correct decision (black solid line). Top-right: Same comparison between "inaccurate" movement trials (red) and trials following an inaccurate movement trial (black). Bottom-right: Same comparison between "unreached" movement trials (red) and trials following an unreached movement trial (black). B. Same as A for trials collected in the CMD task.

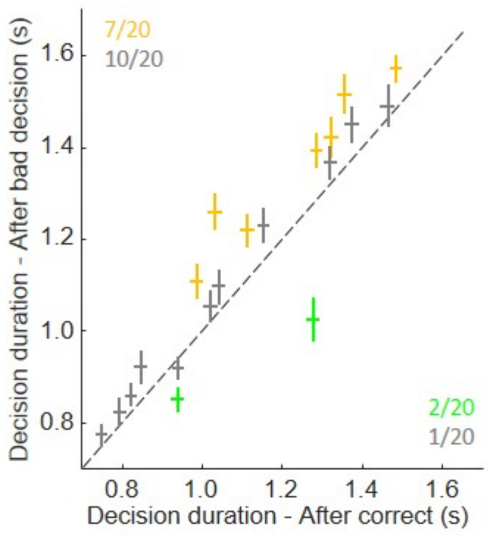


**Supplemental figure 2.** Average decision durations of each subject during post-correct decision trials (x-axis) and post-incorrect decision trials (y-axis) performed in the free-movement duration (FMD) task. Orange (green) pluses indicate the mean and standard error for subjects for whom decision durations following a bad decision were significantly longer (shorter) than decision durations following a correct choice (Wilcoxon-Mann-Whitney test, $p < 0.05$).

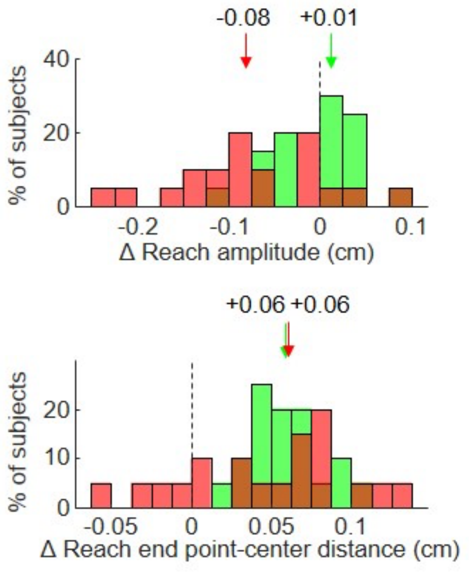


**Supplemental figure 3.** Distribution and comparison of reaching movement amplitude (top) and end-point-center distance (bottom) adjustments depending on the decision outcome in the previous trial (after a correct decision in green and after an error decision in red) of the free-movement duration (FMD) task.

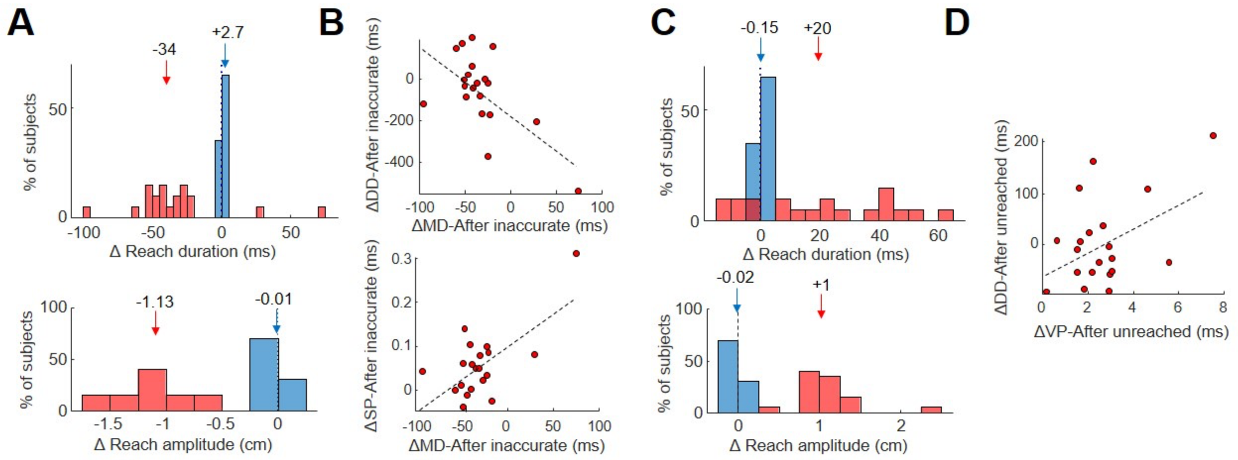


**Supplemental figure 4.** A: Distribution and comparison of subjects' reaching duration (top) and amplitude (bottom) adjustments depending on the movement outcome in the previous trial (after a correct movement in blue and after an inaccurate movement in red) of the free-movement duration (FMD) task. B: Top - Difference of movement duration between inaccurate movement trials and after an inaccurate movement trial for each subject (x-axis) as a function of the difference of decision duration between inaccurate movement trials and after an inaccurate movement trial (y-axis) in the FMD task. The dotted line illustrates a Pearson regression through the data. Bottom - Same as Top for the correlation between movement duration and decision success probability. C. Same as A for unreached movement trials. D. Difference of movement peak velocity between unreached movement trials and after an unreached movement trial for each subject (x-axis) as a function of the difference of decision duration between unreached movement trials and after an unreached movement trial (y-axis) in the FMD task. The dotted line illustrates a Pearson regression through the data.

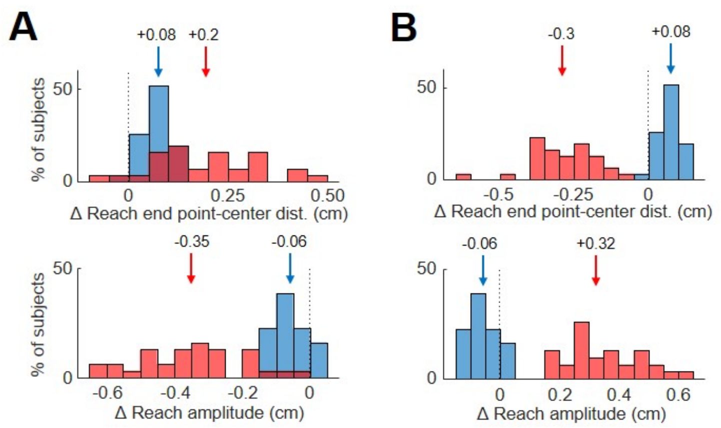


**Supplemental figure 5.** A: Distribution and comparison of subjects' reaching end point-center distance (top) and amplitude (bottom) adjustments depending on the movement outcome in the previous trial (after a correct movement in blue and after an inaccurate movement in red) of the constrained-movement duration (FMD) task. B. Same as A for unreached movement trials.
